# Evaluating and Improving Health Equity and Fairness of Polygenic Scores

**DOI:** 10.1101/2023.09.22.559051

**Authors:** Tianyu Zhang, Lambertus Klei, Peng Liu, Alexandra Chouldechova, Kathryn Roeder, Max G’Sell, Bernie Devlin

## Abstract

Polygenic scores (PGS) are quantitative metrics for predicting phenotypic values, such as human height or disease status. Some PGS methods require only summary statistics of a relevant genome-wide association study (GWAS) for their score. One such method is Lassosum, which inherits the model selection advantages of Lasso to select a meaningful subset of the GWAS single nucleotide polymorphisms as predictors from their association statistics. However, even efficient scores like Lassosum, when derived from European-based GWAS, are poor predictors of phenotype for subjects of non-European ancestry; that is, they have limited portability to other ancestries. To increase the portability of Lassosum, when GWAS information and estimates of linkage disequilibrium are available for both ancestries, we propose Joint-Lassosum. In the simulation settings we explore, Joint-Lassosum provides more accurate PGS compared with other methods, especially when measured in terms of fairness. Like all PGS methods, Joint-Lassosum requires selection of predictors, which are determined by data-driven tuning parameters. We describe a new approach to selecting tuning parameters and note its relevance for model selection for any PGS. We also draw connections to the literature on algorithmic fairness and discuss how Joint-Lassosum can help mitigate fairness-related harms that might result from the use of PGS scores in clinical settings. While no PGS method is likely to be universally portable, due to the diversity of human populations and unequal information content of GWAS for different ancestries, Joint-Lassosum is an effective approach for enhancing portability and reducing predictive bias.

## 1 Introduction

Phenotypic diversity is a hallmark of human populations. When heritable variation underlies within-population diversity, polygenic scores (PGS) are logical, if imperfect, predictors for phenotypic variation [45, 5, 20, 32]. Insofar as we are aware, the concept of genetic scores arose in the animal breeding literature to estimate the breeding values of individuals, usually males [21, 29]. In human genetics, PGS were introduced [35] as a natural byproduct of a genomewide association study (GWAS), from which a subset of SNPs were selected by some threshold (T) for their association with phenotype and some highly dependent SNPs were pruned (P) out to create a sparser set of quasiindependent predictors. Since the original Pruning-and-Thresholding (P&T) method was introduced [35], many and somewhat more accurate methods of PGS have been developed [42, 14, 24, 34, 37, 19, 2, 23, 31, 16, 12]. An ongoing challenge of such PGS, however, is making them portable across ancestries – in other words, if a PGS were derived from a GWAS based on subjects of a certain ancestry, does it provide equivalent prediction for individuals of other ancestries? Disappointingly, the answer is usually no [9, 7, 36, 47, 39]. For example, prediction accuracy for European-based PGS was 4.5 fold lower when applied to individuals of African ancestry [27], raising concerns that clinical use of PGS could exacerbate existing health disparities [7].

In part, migration patterns and limitations to gene flow underlie lack of portability. Populations separated by greater distances or geographic impediments, which tend to hinder gene flow, also tend to cause larger allele frequency differences than obtained for nearby populations. Nonetheless, whenever the distribution of genetic variation within and among populations is assessed, a recurrent theme emerges: variation common in one population is rarely absent in others, even for populations from different continents [38, 4]. Exceptions include variation under natural selection due to environmental forces unique to specific populations. That most common genetic variation is present across diverse ancestries, even if at different frequencies, allows for a simplifying assumption for methods seeking to make PGS more portable: Assume that common variants altering variation of a phenotype in one population (causal variants) have similar but not identical effects in other populations. Another complication is linkage disequilibrium (LD) [18]. Due to extensive LD among proximate SNPs in a population, GWAS does not necessarily reveal causal variants. Rather it reveals associated SNPs, some causal, many neutral but in LD with causal variants, and others falsely associated. Moreover, LD patterns differ among populations, due to human history and resulting allele frequency differences [18]. To overcome this hurdle, it is commonly assumed that LD patterns derived from population resources such as 1000 Genomes, are useful approximations for particular population samples and these can be used to understand patterns of association in different populations.

Some recent reviews [43, 17] delineate the challenges and opportunities in the field, and various methods have been proposed to improve portability [15, 22, 1, 25, 39, 49, 30], including methods such as XPXP[46] and PolyPred[44] that incorporate information beyond GWAS of one phenotype. Building on modern statistical methods, for example, TL-PRS [48] uses transfer learning techniques to improve portability. In this setting, we conjectured that model selection techniques, such as penalized regression embodied in Lassosum [24], could be another useful approach for refining portability. Here we develop Joint-Lassosum, a computationally efficient method to leverage GWAS summary statistics and their proxy LD matrices from two ancestries. This approach is more efficient, computationally, than its natural Bayesian counterpart, PRS-CSx [39]. Using simulations consistent with our assumptions, Joint-Lassosum achieved good prediction performance across various scenarios, including in comparison to prediction from TL-PRS. These simulations use schizophrenia as a model outcome and Europeans and Yorubans as model ancestries. Additionally, we introduce a convenient method for choosing tuning parameters that obviates the need for independent validation data, which is often extremely limited for underrepresented populations. Finally, we draw connections to the literature on algorithmic fairness and assess the fairness of PGS scores across different models. We motivate the disparity in False Discovery Rate (FDR) across groups as a meaningful measure of predictive bias for PGS scores, and show that Joint-Lassosum is effective in reducing this disparity. Furthermore, we demonstrate that underrepresentation is not the only source of disparate predictive performance. Disparities in the predictive performance of PGS scores persist even if Europeans and Yorubans are equally represented in the data. This is because the Yoruban LD structure creates a more challenging estimation problem.

## 2 Materials and Methods

### 2.1 Penalized Linear Regression for PGS

Lassosum [24] is a penalized regression method for derivation of a PGS and requires only GWAS summary statistics and LD information as input. The *l*_1_-penalized regression [40] minimizes the objective function,

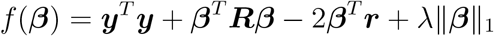

Here ***y*** is the phenotype vector of length *n* (the sample size); ***R*** = ***X***^*T*^ ***X*** is the LD matrix between SNPs (LD block matrix), in the form of a correlation matrix; and ***r*** = ***X***^*T*^ ***y*** is a vector of relationships between the SNPs and phenotype, which can be constructed using GWAS summary statistics. The ∥ ***β*** ∥ _1_ term is the *𝓁*_1_-norm of the SNP effects vector ***β***. Such a sparsity-inducing regularization term shrinks most of the SNP effects to zero, and *λ* is a hyperparameter controlling the regularization strength. GWAS studies usually report phenotype-SNP relationships ***r***, but the SNP-SNP LD levels, which requires the genotypes of the subjects in the study, is often not available. Instead, ***R*** is often estimated based on another resource, such as the 1000 Genomes project. This replacement does not hurt statistical validity, although it could cause the solution to be numerically more sensitive. Therefore, the Lassosum algorithm aims to minimizes the following objective function:

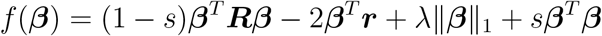

in which *s* is another regularization parameter to ensure a stable solution.

### 2.2 From One Population to Two: Joint-Lassosum

We extend Lassosum by incorporating information from two ancestries. Our proposed method, Joint-Lassosum (JLS, Figure 1), calls for two LD block matrices (***R***_1_, ***R***_2_) and GWAS summary statistics (***r***_1_, ***r***_2_) from two ancestries. These summary statistics can come from different resources. Suppose that the GWAS from population 1 is of large sample size, and therefore its PGS has good performance, whereas we especially wish to improve the PGS for population 2, the “target” population. The (minimization) objective function of Joint-Lassosum is:

**Figure 1.**
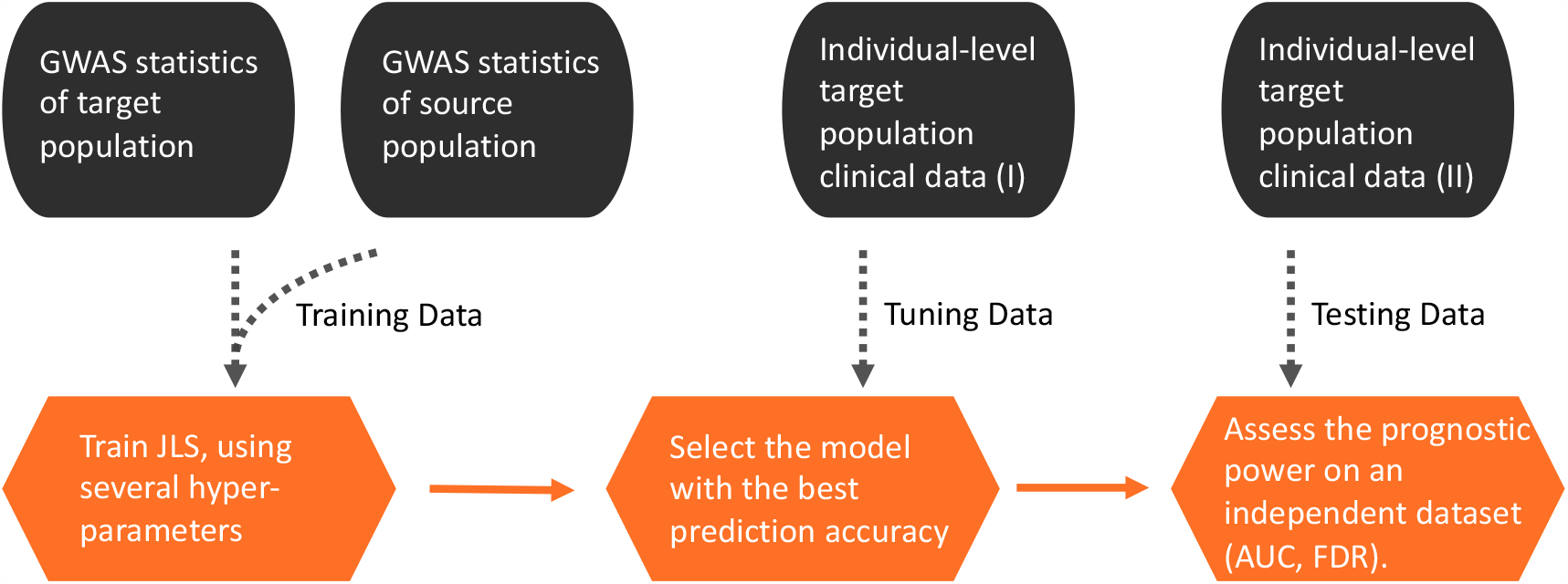
Joint-Lassosum. This flowcharts presents the standard implementation of Joint-Lassosum, which would require independent individual-level target population tuning (for hyper-parameter selection) and testing data (assessing diagnostic accuracy). In practice, the independent Tuning and Testing data sets may not be available, which motivates our synthetic-data based tuning procedure in Section 2.3.

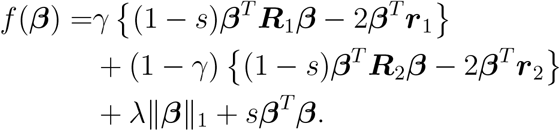

We introduced a new hyperparameter 0 ≤ *γ* ≤ 1 to balance the contribution of the two sets of samples. A larger *γ* up-weights information from population 1, which would have less variability in its association statistics, whereas a smaller *γ* induces a *β* calibrated better to the target population’s genome structure. The objective function can be solved by a coordinate descent algorithm, which is described in detail in Supplement.

There are some direct generalizations of one-population methods to two population problems in the literature. For example, there are methods that take weighted average of two models trained separately for each population (one from the source population and one from the target population). A commonly used method we will call “weighted P&T” [26]; another is “weighted Lassosum” [48]. Although conceptually simple, both methods require tuning models for both populations because the optimal values for parameters will typically be different for each component model. By comparison, Joint-Lassosum only requires tuning two key parameters *γ* and *λ*.

### 2.3 Proposed Parameter Tuning Procedures

Parameter tuning is a common challenge for PGS methods, including all the penalized regression-based PGS methods. In our method, there are three hyperparameters: (1) sparsity penalty parameter *λ*, (2) population weighting parameter *γ*, and (3) shrinkage parameter *s*. The penalty parameter *λ* controls the number of SNPs that have non-zero contribution to the PGS (i.e. the sparsity of the regression coefficients). The weighting parameter *γ* balances the contribution of two populations to the loss function. And the shrinkage parameter *s* is introduced because of numerical concerns. According to our simulation results, the parameter *s* can be trivially set to 0.9. The parameter tuning mainly focuses on the other two more meaningful parameters: sparsity parameter *λ* and population weight parameter *γ*.

To tune these hyperparameters, we propose using a synthetic-sample based procedure. This proposal is inspired by the parametric bootstrap estimation of prediction error described in [10], which is shown to have favorable theoretical guarantees under mild assumptions. The procedure can be briefly described as the following (for a sketch of the framework, see Figure 2):

1. Perform a preliminary fitting of Joint-Lassosum under a set of candidate hyperparameters. Each candidate hyperparameter corresponds to one candidate model.
2. Use one of the fitted coefficients 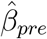 and publicly available genotype information to generate (individual-level) synthetic samples. They are first generated as individual-level samples. The synthetic sample should have a similar sample size, noise level and study design as the original data set of interest. (Note that this information can be obtained by examining summary statistics from the GWAS samples).
3. We employ standard (cross-)validation techniques with the synthetic samples to select the hyperparameters. This is possible because we have access to the genotypes of each synthetic individual and their phenotype. More specifically, this step can be further divided as:
  (3.1) Split the data generated in step (2) into training and testing sets.
  (3.2) Fit models using the training set, with the same candidate hyperparameters as in step (1).
  (3.3) Use the testing set to evaluate the performance, determine the better candidate hyperparameter. A range of sparsity hyperparameters (*λ*s) may produce similar performance (AUCs). In this situation, we recommend choosing the more regularized model (i.e., the model that selects fewer non-zero regression coefficients).
4. Choose the model, among those created in step 1, that corresponds to the hyper-parameter selected in step 3.

**Figure 2.**
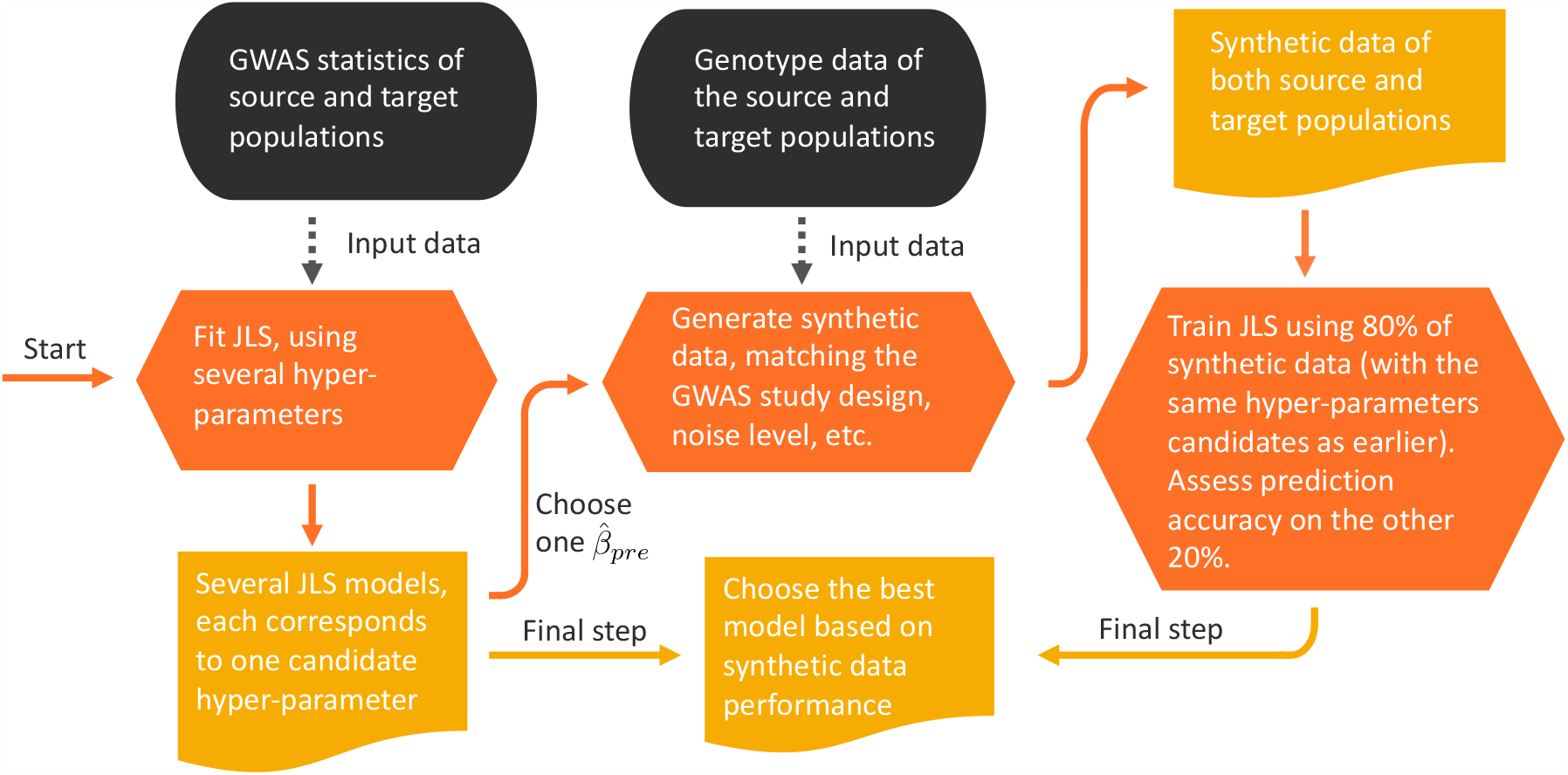
Framework of synthetic data-based hyperparameter tuning procedure, taking the proposed Joint-Lassosum method as an example. In this work, the “source population” usually refers to subjects of European ancestry and the “target population” are those of African ancestry.

This procedure is general and is applicable to other methods as well.

### 2.4 Simulations using realistic linkage disequilibrium data

We performed simulations to evaluate the performance of Joint-Lassosum under various scenarios and compare its efficiency with some other PGS algorithms. Simulations were based on two populations, one of European and another of African ancestry. Imitating the current-day reality, for most simulations the sample size was larger for the sample of European-ancestry and we wish to transfer what is learned from the larger European to the smaller African population. To generate genotypes, we used data on inferred haplotypes from individuals who contributed DNA to the 1000 Genomes project, specifically haplotypes from 179 individuals of European (CEU) and 178 African (Yoruban, YRI) ancestry. From these data, we selected SNPs that were haplotyped across all samples; had minor allele frequency greater than 0.01 and p-values greater than 0.005 from the two tests of Hardy-Weinberg Equilibrium, one for each ancestry group; were autosomal; and fell into one of the 2,559 and 1,681 linkage disequilibrium (LD) blocks for YRI and CEU, respectively, as defined by [3] . In total, 5,630,745 SNPs met these criteria.

Within each ancestry, genotypes for each of N individuals were generated by LD block (e.g., within a block, sample from the 356 haplotypes that were available for YRI). For each of the N individuals, we randomly selected two of the haplotypes with the restriction that they could not be the same. Performing this operation over all haplotype blocks creates the combined set of genotypes for 5,630,745 SNPs; among these SNPs, the LD structure of the original CEU and YRI samples were largely preserved, as were the relative allele frequencies.

From these SNPs, we randomly selected 4000 causal variants after meeting these conditions: SNPs were chosen based on the relative length of each chromosome versus their total length; the first causal SNP was chosen to be between 100Kb and 250Kb from the first SNP on the chromosome; the remaining causal SNPs were then spread approximately equidistant across the chromosome. Average distance between causal SNPs was approximately 690Kb.

Our next goal was to obtain the binary phenotypes for individuals using the standard threshold model from quantitative genetics [11], which assumes an underlying normal distribution for population liability for a binary trait, and under which an individual is affected if their liability exceeds a threshold *t*. The prevalence of affection status in the population, which we set to 1.5% for our simulations, determined *t*. To generate causal effects, we fixed the variance due to environmental effects to 1, which is typical for the threshold model. The total variance due to genetics effect was then *σ*_*g*_ = (1 − *h*^2^)*/h*^2^, in which *h*^2^ is the heritability of a binary trait on the liability scale. After distributing the genetic variance evenly across loci, *τ* = *σ*_*g*_*/*4, 000, the effect size for each SNP *i* was defined as (*τ/*(2*p*_*i*_*q*_*i*_))^1*/*2^, where *p*_*i*_ is the relative frequency of causal allele *i* in the population sample and *q*_*i*_ = 1 − *p*_*i*_.

The genetic score for each simulated subject was determined by summing the number of risk alleles times the effect size at each locus. Due to finite size of subjects and causal variants, we adjusted these genetic scores so that their mean is zero. To account for environmental effects on the phenotype, a random number drawn from *N* (0, 1) was added to the individual’s genetic score to obtain their phenotypic score on the liability scale. Affection status was then assigned based on whether the phenotypic score exceed *t* from the normal distribution *N* (0, *σ*_*g*_ + 1). Rejection sampling was performed until *N/*2 affected and *N/*2 unaffected individuals were obtained. For each simulation we created a population used to calculate the GWAS (TRN), tuning parameters (TUNE), and testing (TST) for both CEU and YRI. Unless noted otherwise, for each of the TRN, TUNE or TST, we generated 20,000 CEU subjects and 4,000 YRI subjects. To obtain references populations for quantifying LD among SNPs, we generated random sets of subjects of CEU and YRI ancestry with the same size. For these samples, affection status was irrelevant.

## 3 Results

### 3.1 Performance of PGS

We investigate performance of some one- and two-population PGS methods under various simulated scenarios. In the first scenario, we set the heritability of liability of the phenotype to 80%, and set the genetic variance explained by each causal variant to be the same over SNPs and populations (CEU and YRI). Because relative allele frequencies of SNPs typically differ between populations, the realized effects of the SNPs on phenotype differed as well. By contrast to a one-population method (Lassosum, LS), two-population PGS algorithms substantially improve prediction of affection status for the smaller YRI sample (Figure 3A). Of the two-population PGS algorithms, Joint-Lassosum (JLS) slightly outperforms Transfer Learning (TL, [48]) and weighted Lassosum (WLS). Similar patterns were obtained for lower heritablity (50%), although prediction accuracy is correspondingly diminished over all methods too (Figure S1A). The same patterns hold when there are different heritabilities, 80% for CEU and 60% for YRI populations (Figure S1B).

**Figure 3.**
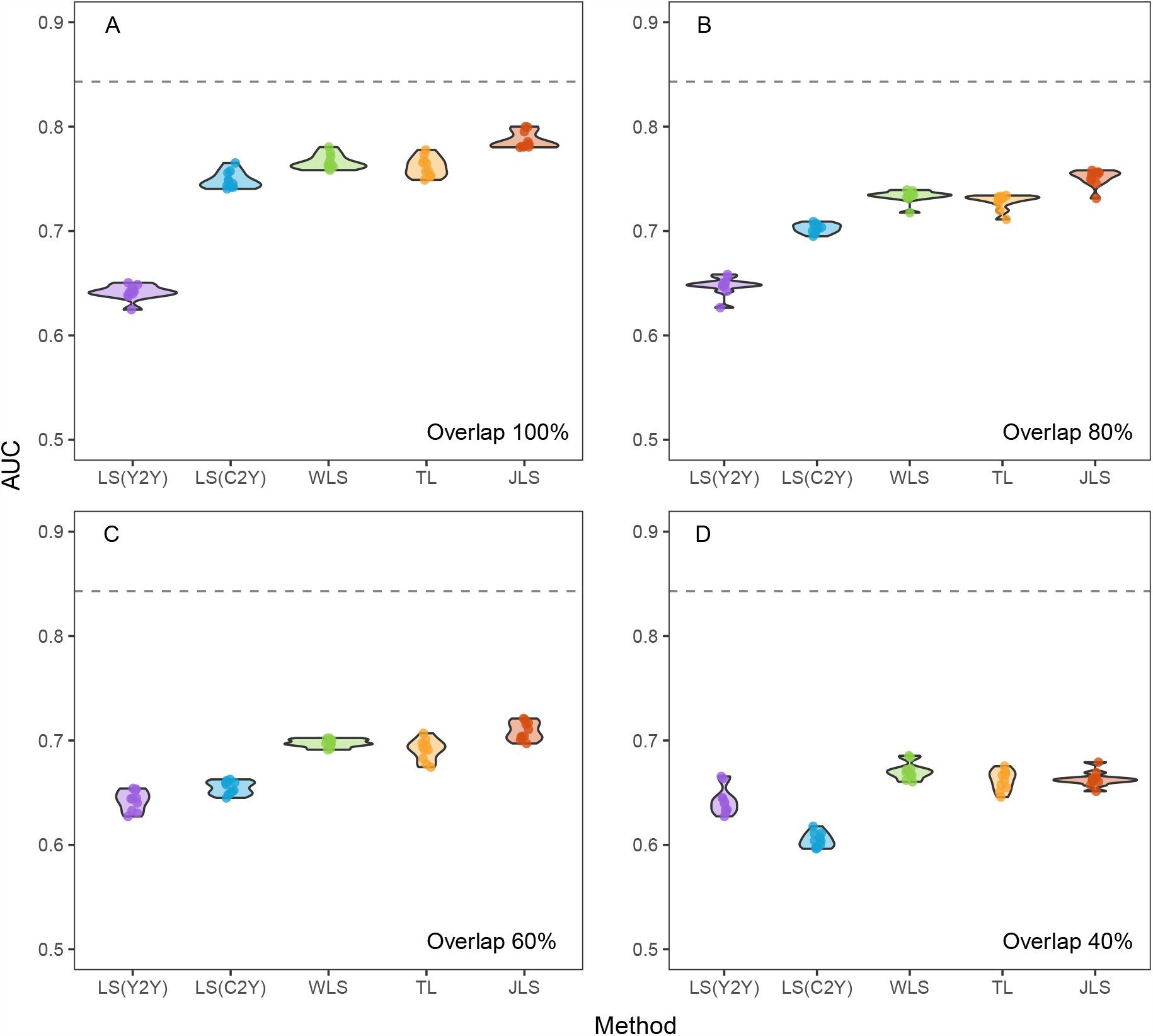
Prediction accuracy of PGS algorithms, as measured by Area Under the receiver operating characteristic Curve (AUC) **A**. The dashed line is the AUC of a Lassoum (LS) PGS for a CEU sample, trained using another sample from the CEU population. Also shown are a LS PGS for a Yoruban (YRI) sample, trained using another sample from the YRI LS(Y2Y) or CEU LS(C2Y) population; a PGS obtained as a weighted combination of the Y2Y and C2Y PGS, denoted as WLS; a PGS obtained by a transfer learning method (TL); and a PGS obtained by Joint Lassosum (JLS). In **B**-**D**, results are shown for diminishing levels of genetic correlation (0.80, **B**; 0.60, **C**; and 0.4, **D**) between the trait in the CEU and YRI populations.

The meaning of prediction accuracy, as summarized by AUC, is arguably obscure for clinical practice. Suppose clinicians wanted to apply a preventative treatment to adolescents who, without it, would go on to be diagnosed with schizophrenia. To identify those at greatest risk, they selected individuals with extreme PGS scores, say, exceeding 2. The fraction of individuals selected who would never be affected, the false discovery rate (FDR), differs markedly across populations and methods (Figure 4A). For the larger CEU population, when trained with another CEU samples, only 3.4% of the individuals would be falsely treated, whereas a much larger fraction of YRI would be, regardless of method. As expected based on AUC (Figure 3A), JLS performs slightly better than TL at minimizing the FDR (7.5% versus 8.2%), and far better than LS(Y2Y) (21.2%), although none performs as well as LS(C2C). As expected, more extreme results were obtained for lower (Figure S2A) or unequal heritabilities (Figure S2B).

**Figure 4.**
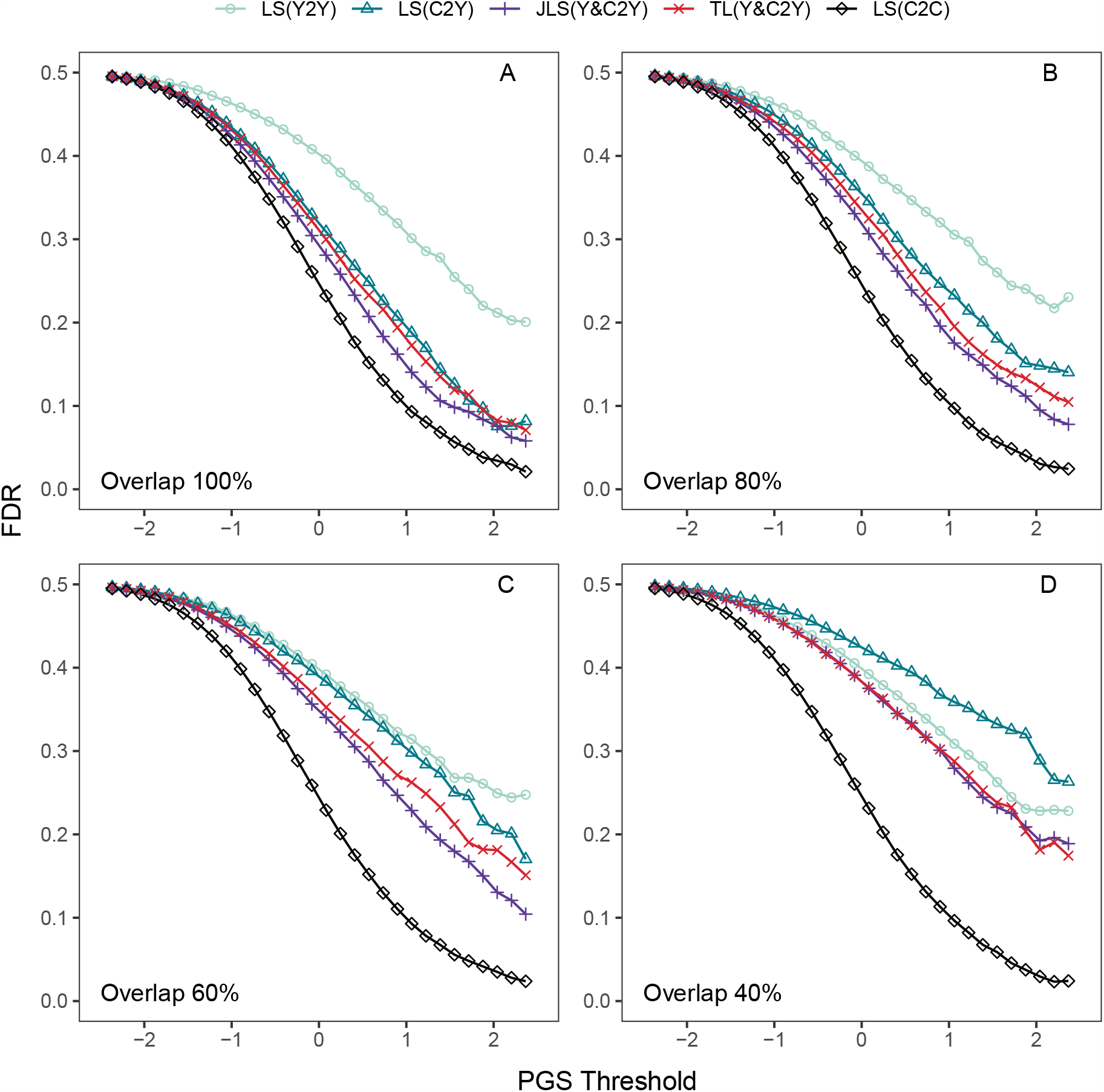
Prediction inaccuracy, as highlighted by the false discovery rate (FDR). The legend uses the format Method(training population + 2(to) + testing population) population, as described in Figure 3.

In the first simulated scenario (Figure 3A), the genetic correlation of the trait over the two populations was 1.0, even though the effects of risk alleles differed between the populations. We also explored prediction performance when the heritability of the trait was held constant over populations, but the genetic correlation was less than unity. We explored three levels of genetic correlation, namely 0.8, 0.6, and 0.4 (Figure 3B-D, Figure 4B-D), which were accomplished by reducing the overlap of causal loci (i.e., 80, 60, and 40% shared causal SNPs respectively). As the genetic correlation declines, the prediction accuracy declines for all of the two-population methods. Nonetheless they always outperformed the one-population methods. Notably, despite the fivefold smaller size of the YRI sample, versus CEU, as the genetic correlation fell to 0.4, the predictive accuracy of the PGS for YRI trained on YRI outperformed that of YRI scored by CEU (Figure 3D). Paralleling results for AUC, FDR increased with diminishing genetic correlation (Figure 4B-D). If clinicians were to use a PGS trained on CEU samples for prediction of YRI risk, false treatment rates would be very high, approaching 30%.

To explore the effect of the size of the YRI sample, we generated 20,000 YRI under the 60% overlap scenario and compared the FDR over a range of PGS thresholds for four combinations of method, training population, and testing population. Even when YRI and CEU are equally represented in the data (Figure 5A), the FDR for LS(Y2Y) remains significantly greater than that of LS(C2C) at all thresholds. This suggests that shorter LD in YRI makes the estimation process inherently more challenging. Nonetheless, the poor performance of PGS for the YRI population, as seen in most of our results, is largely due to smaller sample size. Furthermore, JLS improves performance over LS for both YRI and CEU populations, even when the training sample sizes are both large (Figure 5A). This suggests that any bias induced by using heterogenous training data is overcome by the reduced variance of the larger sample size. Next, by subsampling the 20,000 YRI to obtain a range of sample sizes from 4,000 to 20,000, we varied the proportion of the YRI to CEU population from 0.2 to 1.0 and evaluated FDR for YRI at a threshold of 2 (Figure 5B). Notably, while LS is always inferior to JLS, the relative improvement produced by JLS declines as the proportion grows.

**Figure 5.**
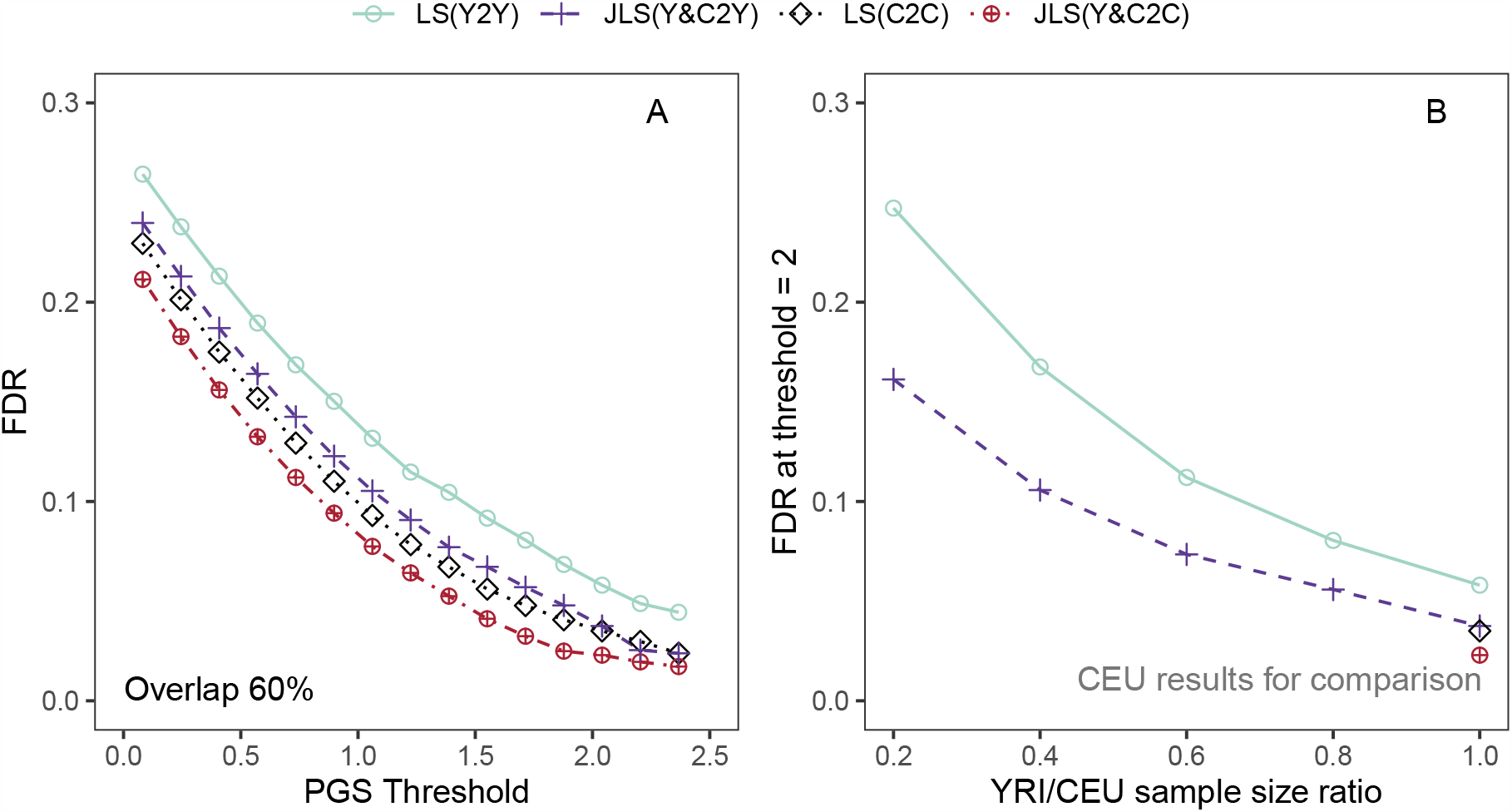
Prediction inaccuracy, as highlighted by the false discovery rate (FDR). The legend uses the format Method(training population + 2(to) + testing population) population, as described in Figure 3A. (A) Large, equal sample size setting (YRI = CEU = 20,000). (B) PGS threshold is set to be 2, varying YRI sample size (from 4,000 to 20,000). The data points at (x-axis =)2 in panel A are identical to those at 1 in panel B.

### 3.2 Parameter Tuning with Synthetic Samples

While hyperparameter selection is critical for all PGS algorithms, an independent data set for such tuning is often unavailable, especially for underrepresented populations. In Methods, we presented a synthetic-tuning approach to overcome this challenge. Here, through simulations, we evaluated the proposed approach. In these simulations, we generated training and testing samples (CEU and YRI) and calculated summary GWAS statistics. Using these summary statistics, multiple Joint-Lassosum models were fitted under 30 different combinations of hyper-parameters *γ, λ* (three choices of *γ* times ten choices of *λ*). These 30 models all had different prediction accuracy, based on the test data, and we defined the best of them as the “Oracle (testing) AUC”. We performed 10 such simulations, so there were 10 “Oracle AUC” for each population.

To perform the proposed model selection procedure described in Section 2.3, we used one of the 30 models’ regression coefficients 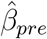 to generate synthetic data. (In practice, it is up to the user which 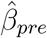 is implemented because no other reference information is available.) We chose four different 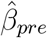 to generate the synthetic data: (I) *γ* = 0.8, *λ* = 7 ×10^*−*3^; (II) *γ* = 0.5, *λ* = 7 ×10^*−*3^; (III) *γ* = 0.2, *λ* = 1.5 ×10^*−*2^ and (IV) *γ* = 0.8, *λ* = 2.5 ×10^*−*3^. The Oracle testing AUC in general is achieved around *γ* = 0.8, *λ* = 2 ×10^*−*2^, which we purposely avoided in the 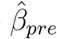 settings. Using these synthetic tuning data, the hyperparameters yielding best prediction from the synthetic test data are chosen. The Joint-Lassosum model corresponding to these hyperparameters, one of the 30 models, is then applied to the true target data to determine the realized AUC, which is then compared to the corresponding Oracle (Figure 6). For comparison, we also generated independent tuning data, 20,000 CEU and 4,000 YRI samples, and used them to perform model selection with cross-validation (CV). In each repeat, cross-validation identified one model that is expected to perform best and the selected models’ testing AUC was contrasted to the corresponding Oracle (Figure 6). Leveraging extra individual-level information, CV can more consistently select models of almost the best performance. However, synthetic-data model selection can still help the investigators to effectively exclude severely under-performing ones without requiring any (extra) individual-level information. Typically, more than half of the 30 candidate models have YRI testing AUC *<* 0.7 (the worst can be as low as 0.6), and we observe that the proposed procedure is likely to select models of AUC between 0.75 and 0.8 (as presented in the figure)—Synthetic (IV) is more under-performing, usually getting AUC around 0.75.

**Figure 6.**
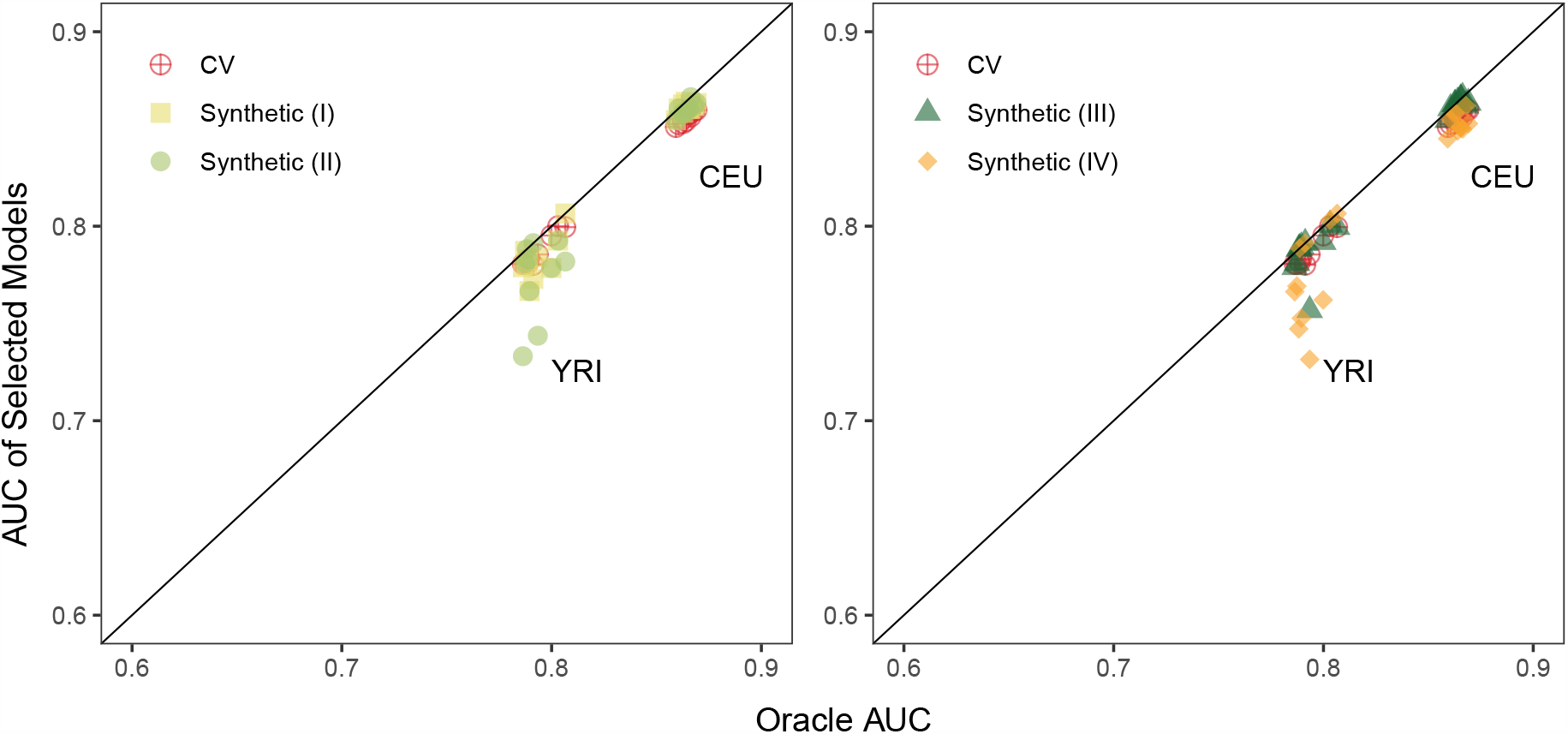
Synthetic data-based tuning method gives comparable results to cross-validation (CV) and the Oracle for two populations, CEU and YRI. The Oracle testing AUC on the *x*-axis is the best AUC that can be achieved by the candidate hyperparameters. Synthetic data sets are generated under four different choices of 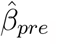 which are estimated under different hyperparameter combinations.

## 4 Discussion

Polygenic scores are typically developed from one major ancestry group, such as European ancestry. Because of human diversity, unequal information content of GWAS for different ancestries, and LD among nearby SNPs, such PGS have limited portability to other ancestries. Here we develop a portable PGS that requires only summary statistics from a relevant genome-wide association study (GWAS), and has the model selection advantages of the well-characterized Lasso method. We present Joint-Lassosum (JLS) (Figure 1), which integrates information for two populations to increase portability of the PGS from the larger “source” population to the smaller “target” population. Simulations show JLS provides portability comparable with other methods (Figures 3, 4, 5). JLS requires selection of data-driven tuning parameters, as do all PGS methods. We describe an approach to tuning that obviates the need for additional population samples, beyond the original GWAS, by generating synthetic population samples for tuning from the GWAS results (Figures 2, 6). JLS, and our proposed synthetic tuning, are effective approaches for enhancing PGS portability. In large part our results emphasize the False Discovery Rate (FDR) as a measure of performance, in contrast to AUC, a popular metric. This is because AUC can fail to differentiate performance of PGS methods in a meaningful way, while FDR provides a metric that is meaningful in practice.

We can also view the results of the simulations used to evaluate JLS (Figures 4, 5) through the lens of fairness-related harms [8]. Specifically, we are concerned with the potential for *allocative harms*, wherein a resource or burden is inequitably distributed due to the differential performance of a model across groups (here, ancestry groups). To ground the discussion, consider the hypothetical clinical screening scenario described for our simulations. Herein, a PGS model is to be used to identify adolescents who are at greatest risk of schizophrenia, for the purpose of providing therapeutic treatment aimed at prevention. In this setting, allocative harms can arise in two ways: (i) failing to treat individuals who develop schizophrenia; and (ii) unnecessarily treating individuals who would not develop schizophrenia even in the absence of treatment. We focus here on the second setting—over-treatment due to false positive predictions.

Individuals who are flagged in adolescence as being high risk for developing schizophrenia and subsequently treated would be subject to at least two sources of harm. First, beyond burdens such as cost, any treatment could have significant side effects, and side effects of drugs designed to prevent psychosis are numerous [28]. Second, such individuals could unnecessarily and adversely alter their lives going forward due to perceived risk. We can assess the extent to which a given PGS model would inequitably subject individuals to these harms by comparing the FDR across groups. Because FDR equals one minus the positive predictive value, comparing FDR is equivalent to assessing a model for *predictive parity* —that is, equivalence of positive predictive values—a fairness metric considered in risk assessment contexts such as criminal justice and health care [6, 41, 33]. For instance, consider results in Figure 4C, in which there is 60% overlap of risk variants in the Yoruban and CEU-European populations. We can see from that, in the absence of joint modeling, FDR rates at all score thresholds are considerably lower for the CEU-European population (LS(C2C)) compared to the Yoruban population (LS(Y2Y)). Joint modeling helps to bridge this gap, with the proposed JLS method producing the greatest decrease in FDR. For instance at a threshold of 2, which is high enough to be plausibly clinically relevant, the FDR for YRI drops from 25% to 13% when applying JLS. This brings the FDR disparity from 25% -3% = 22% to 13% -3% = 10%. Thus, while disparity persists, it is greatly mitigated by applying JLS. For simplicity, we focused our analyses on two continental ancestry groups. Nonetheless, the results have direct relevance to African Americans, whose ancestry typically is a mixture of African and European ancestries. Clearly the fairness discrepancies arising from unequal sample sizes for GWAS, as seen in Figures 4, 5, apply with equal force to African Americans, who are typically underrepresented in genetic studies. Moreover, of relevance to our hypothetical clinical screening scenario, African Americans also tend to be disproportionately diagnosed with psychotic disorders, at least in part due to diagnostic bias [13]. This underscores the importance of ensuring accuracy of PGS for minority populations, if PGS is to be used in a clinical setting, as a matter of fairness and to minimize harms.

There are now a variety of two-population methods for enhancing portability [15, 22, 1, 25, 39, 49]. Undoubtedly some of these methods will yield better prediction than JLS for some pairs of populations and samples. Nonetheless, whenever multiple analytical methods have different advantages in different settings, ensembles of these methods, appropriately tuned, will perform at least as well as and usually better than any one of the methods. Hence an ensemble of current two-population methods is a viable option for greater prediction accuracy.

## Acknowledgements

This project was funded by National Institute of Mental Health (NIMH) grants R01MH123184 and R37MH057881.

