## Supplementary Materials for "Evaluating and Improving Health Equity and Fairness of Polygenic Scores"

September 22, 2023

#### 1 Details of the methods

##### 1.1 Fitting Joint-Lassosum Estimators

To simplify the notation,we will use $y$ for $y_1$, $y'$ for $y_2$, $X$ for $X_1$, $X'$ for $X_2$, etc. We consider $\boldsymbol{y}$ and $\boldsymbol{y'}$ are phenotype vectors, and $\boldsymbol{X}$ and $\boldsymbol{X'}$ are standardized design matrices of genotype for the two ancestries respectively. The objective function of xxx can be specified as follows:

$$
\begin{aligned}
f(\boldsymbol{\beta}) =& \gamma \left(\boldsymbol{y}^T\boldsymbol{y} + (1-s)\boldsymbol{\beta}^T\boldsymbol{R}\boldsymbol{\beta} - 2\boldsymbol{\beta}^T\boldsymbol{r}\right) \\
&+ (1-\gamma)\left(\boldsymbol{y'}^T\boldsymbol{y'} + (1-s)\boldsymbol{\beta}^T\boldsymbol{R'}\boldsymbol{\beta} - 2\boldsymbol{\beta}^T\boldsymbol{r'}\right) \\
&+ s\boldsymbol{\beta}^T\boldsymbol{\beta} + 2\sum_j \lambda|\beta_j|
\end{aligned}
$$

where $\lambda$ is the penalty tuning parameter controlling the number of nonzero elements in $\boldsymbol{\beta}$. $\gamma \in [0,1]$ is a tuning parameter controlling the balance between the two ancestries, and $s$ is a shrinkage parameter for ridge penalty as specified in lassosum.

Let $\tilde{\boldsymbol{X}} = \sqrt{1-s}\boldsymbol{X}_r$, the above objective function can be written as:

$$\begin{aligned} f(\boldsymbol{\beta}) &= \gamma \left( \boldsymbol{y}^T \boldsymbol{y} + \boldsymbol{\beta}^T \tilde{\boldsymbol{X}}^T \tilde{\boldsymbol{X}} \boldsymbol{\beta} - 2\boldsymbol{\beta}^T \boldsymbol{r} + s\boldsymbol{\beta}^T \boldsymbol{\beta} + 2\sum_j \lambda |\beta_j| \right) + \\ & \quad (1-\gamma) \left( \boldsymbol{y'}^T \boldsymbol{y'} + \boldsymbol{\beta}^T \tilde{\boldsymbol{X}}'^T \tilde{\boldsymbol{X}}' \boldsymbol{\beta} - 2\boldsymbol{\beta}^T \boldsymbol{r'} + s\boldsymbol{\beta}^T \boldsymbol{\beta} + 2\sum_j \lambda |\beta_j| \right). \\ &= \gamma f_1(\boldsymbol{\beta}) + (1-\gamma) f_2(\boldsymbol{\beta}) \end{aligned}$$

For SNP $j$, the objective function can be simplified as:

$$f(\boldsymbol{\beta}_j) = \gamma f_1(\boldsymbol{\beta}_j) + (1-\gamma) f_2(\boldsymbol{\beta}_j),$$

where $f_1(\boldsymbol{\beta}_j) = \beta_j^2 \left( \tilde{\boldsymbol{X}}_j^T \tilde{\boldsymbol{X}}_j + s \right) - 2\beta_j \left( r_j - \tilde{\boldsymbol{X}}_j^T \tilde{\boldsymbol{X}}_{-j} \boldsymbol{\beta}_{-j} \right) + 2\lambda |\beta_j|$ and $f_2(\boldsymbol{\beta}_j) = \beta_j^2 \left( \tilde{\boldsymbol{X}}'^T_j \tilde{\boldsymbol{X}}'_j + s \right) - 2\beta_j \left( r'_j - \tilde{\boldsymbol{X}}'^T_j \tilde{\boldsymbol{X}}'_{-j} \boldsymbol{\beta}_{-j} \right) + 2\lambda |\beta_j|$

Consider a coordinate descent step for solving the above equation. It's reasonable to assume SNPs from different LD blocks are not correlated. For computational efficiency, we would like to compute the gradient at $\beta_i$ by its corresponding LD blocks. Specifically, suppose SNP $i$ is in $j^{th}$ LD block in population 1 and $k^{th}$ LD block in population 2:

$$f_1(\boldsymbol{\beta}_j) = \beta_j^2 \left( \tilde{\boldsymbol{X}}_j^T \tilde{\boldsymbol{X}}_j + s \right) - 2\beta_j \left( r_j - \tilde{\boldsymbol{X}}_j^T \tilde{\boldsymbol{X}}_{-j}^{[k]} \boldsymbol{\beta}_{-j}^{[k]} \right) + 2\lambda |\beta_j|$$

$$f_2(\boldsymbol{\beta}_j) = \beta_j^2 \left( \tilde{\boldsymbol{X}}'^T_j \tilde{\boldsymbol{X}}'_j + s \right) - 2\beta_j \left( r'_j - \tilde{\boldsymbol{X}}'^T_j \tilde{\boldsymbol{X}}'^{[k']}_{-j} \boldsymbol{\beta}_{-j}^{[k']} \right) + 2\lambda |\beta_j|$$

where $\tilde{\boldsymbol{X}}_{-j}^{[k]}$ is the submatrix for LD block $k$ excluding $\tilde{\boldsymbol{X}}_j$, and $\boldsymbol{\beta}_{-j}^{[k]}$ is the subvector for LD block $k$ excluding $\beta_j$.

Conditional on $\boldsymbol{\beta} = \tilde{\boldsymbol{\beta}}$, we compute gradient at $\beta_j$.

- When $\beta_j > 0$:

$$\left. \frac{\partial f_1(\beta_j)}{\partial \beta_j} \right|_{\beta^{[k]} = \tilde{\beta}^{[k]}} = 2\beta_j \left( \tilde{\boldsymbol{X}}_j^T \tilde{\boldsymbol{X}}_j + s \right) - 2 \left( r_j - \tilde{\boldsymbol{X}}_j^T \tilde{\boldsymbol{X}}_{-j}^{[k]} \tilde{\boldsymbol{\beta}}_{-j}^{[k]} \right) + 2\lambda$$

  Likewise we can compute the gradient for $f_2(\beta_j)$. Setting $\frac{\partial f(\beta_j)}{\partial \beta_j} = 0$, after simple transformation, the coodinate-wise update for $\beta_j > 0$ has the form:

$$\beta_j = \begin{cases} 0 & \text{otherwise} \\ \frac{u_j - \lambda}{\gamma \tilde{\boldsymbol{X}}_j^T \tilde{\boldsymbol{X}}_j + (1-\gamma) \tilde{\boldsymbol{X}}'^T_j \tilde{\boldsymbol{X}}'_j + s} & \text{if } u_j - \lambda > 0 \end{cases}$$

where $u_j = \gamma \left( r_j - \tilde{\boldsymbol{X}}_j^T \tilde{\boldsymbol{X}}^{[k]}_{-j} \tilde{\boldsymbol{\beta}}^{[k]}_{-j} \right) + (1-\gamma) \left( r'_j - \tilde{\boldsymbol{X}}'^T_j \tilde{\boldsymbol{X}}'^{[k']}_{-j} \tilde{\boldsymbol{\beta}}^{[k']}_{-j} \right)$

- When $\beta_j < 0$: Similarly, the coodinate-wise update for $\beta_j < 0$ has the form:

$$\beta_j = \begin{cases} 0 & \text{otherwise} \\ \frac{u_j + \lambda}{\gamma \tilde{\boldsymbol{X}}_j^T \tilde{\boldsymbol{X}}_j + (1-\gamma) \tilde{\boldsymbol{X}}'^T_j \tilde{\boldsymbol{X}}'_j + s} & \text{if } u_j + \lambda < 0 \end{cases}$$

where $u_j = \gamma \left( r_j - \tilde{\boldsymbol{X}}_j^T \tilde{\boldsymbol{X}}^{[j]}_{-j} \tilde{\boldsymbol{\beta}}^{[k]}_{-j} \right) + (1-\gamma) \left( r'_j - \tilde{\boldsymbol{X}}'^T_j \tilde{\boldsymbol{X}}'^{[k]}_{-j} \tilde{\boldsymbol{\beta}}^{[k]}_{-j} \right)$

Combining the above two cases, the coodinate-wise update for $\beta_j$ is

$$\beta_j = \begin{cases} 0 & \text{otherwise} \\ \frac{\text{sign}(u_j)(|u_j| - \lambda)}{\gamma \tilde{\boldsymbol{X}}_j^T \tilde{\boldsymbol{X}}_j + (1-\gamma) \tilde{\boldsymbol{X}}'^T_j \tilde{\boldsymbol{X}}'_j + s} & \text{if } |u_j| > \lambda \end{cases} \tag{1}$$

The running time of our software is roughly linear with respect to the number of SNPs. For GWAS of 500,000 SNPs, using 10,000 samples to calculate the correlation matrices, fitting one single Joint-Lassosum model typically takes less than 60 minutes.

#### 1.2 Normalization Details of the Data

In the main manuscript, we discussed 1-norm (i.e. $\|\beta\|_1 = \sum_{i=1}^p |\beta_i|$) penalized regression using summary statistics. For clarification and proper application of our proposed methods, we include a short note about the normalization procedure of the data. Recall that in the simplest one population case, penalized regression for PGSs aims to find $\beta$ that minimizes the following objective function:

$$f(\beta) = (1-s)\beta^\top R \beta - 2\beta^\top r + \lambda \|\beta\|_1 + s\beta^\top \beta,$$

with $R = X^\top X$ and $r = X^\top y$. The genotype matrix $X$ is of dimension $n \times p$ ($n$ is sample size, $p$ is the number of SNPs) and we used $y$ to denote the phenotype vector ($y = 1$ for case and $y = 0$ for control), which is a $n$-length vector.

From GWAS statistics, we can calculate the sample correlation coefficient (covariance divided by product of standard deviations) between each SNP and the phenotype. However, it is only numerically equal to the desired quantity $r = X^\top y$ after under certaion normalization. Specifically, each column of $X$ should be normalized to have mean 0 and unit 2-norm (i.e., $\sum_{i=1}^n x_{ij}^2 = 1$ for each $j = 1, 2, ..., p$, $x_{ij}$ is the $(i, j)$-th element of matrix $X$). Similarly, $y$ also needs to have zero mean and unit 2-norm ($\sum_{i=1}^n y_i^2 = 1$).

In the two-population case, we assume the same normalization *for each* $X_1, y_1$ and $X_2, y_2$. This makes the two ancestries of different sample size numerically on a comparable scale. The above normalization details become more relevant when investigators applying synthetic-data based parameter tuning. When calculating the PGS for each subject or the calibration slope (Section 3), we recommend the users double-check proper normalization has been applied.

### 2 Supplementary Simulation Results

#### 2.1 Lower heritability setting

In Figure S1 and Figure S2, we present the prediction AUC and FDR results for the "lower-heritability" setting. For discussion and comparison with the other settings, see Section 3 in the main manuscript.

#### 2.2 Equal large sample size setting

We also examined the performance of several lasso-type methods when there are equal number of samples simulated from the YRI and CEU populations ($20,000$ each). In Figure S3, we report the testing AUC on the YRI population in the left panel and that of the CEU population in the right panel. As expected, the one-population Lassosum models performs better when the training and testing populations are the same (Y2Y and C2C). Both populations would further benefit from two-population methods since they would effective increase the sample size. YRI population in general has a lower testing AUC than the CEU population. We also report FDR results in Figure S4.

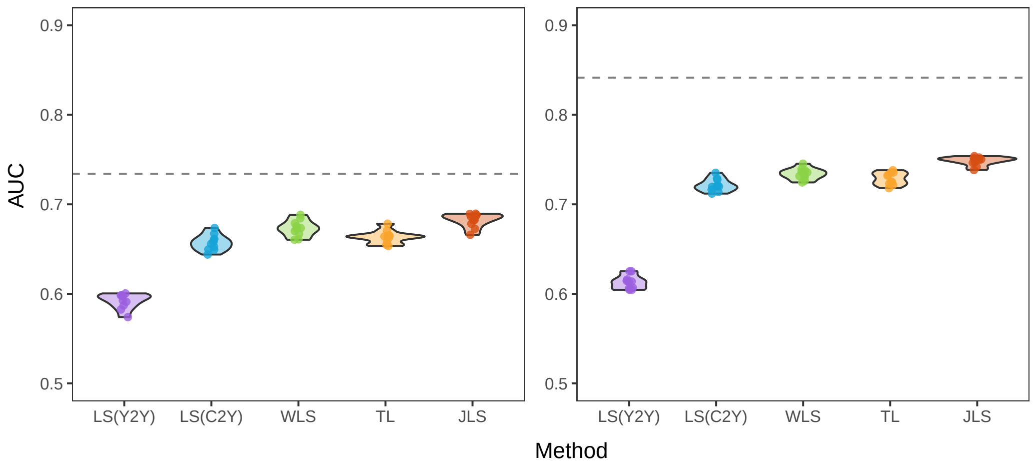


Figure S1: Prediction AUC under lower heritability. Left, Low heritability setting $h^2 = 0.5$. Overlap percentage is 100%; Right, $h^2 = 0.6$ for YRI and 0.8 for CEU.

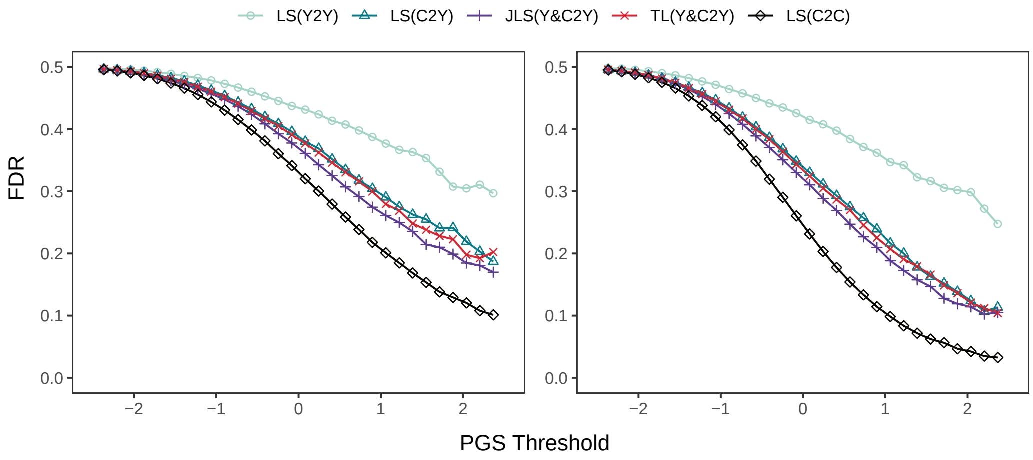


Figure S2: FDR under lower heritability. Left, $h^2 = 0.5$; Right, $h^2 = 0.6$ for YRI and 0.8 for CEU.

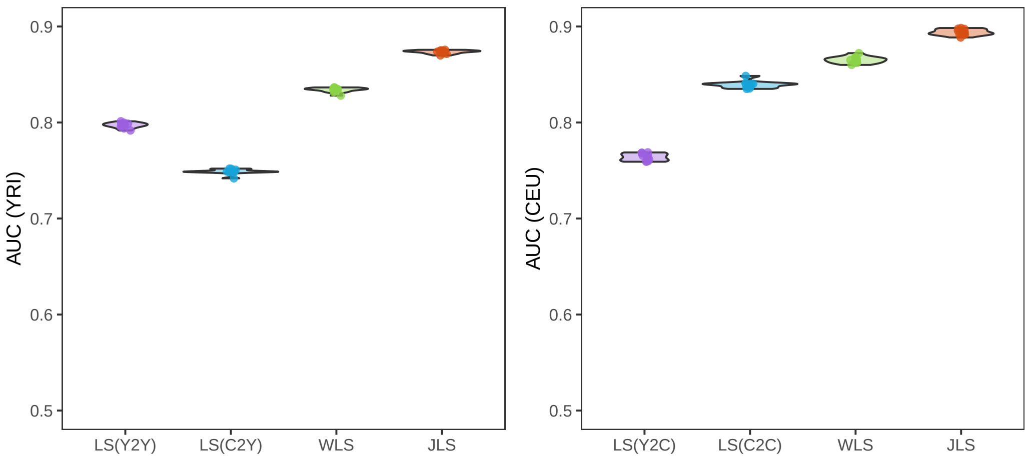


Figure S3: AUC, equal sample size setting.

### 3 Hyperparameter Selection without Tuning Samples

In this section, we give a more formal presentation of the our proposed hyperparameter strategy when there is only a training sample with summary statistics available.

We present more details of the procedure in Algorithm 1. To better match the GWAS study design, which would improve the quality of the selected hyperparameter, we need to "calibrate" (perform a linear transformation) the predicted PGS for each subject before drawing their phenotype label. Below we present a sketch of how we arrived at our calibration formula.

For each subject $i \in \{1, 2, ..., n\}$, we use $X_i \in \mathbb{R}^p$ to denote their genotype vector. We use $\hat{\beta}_{pre} \in \mathbb{R}^p$ to denote the preliminary model we use to generate synthetic data. Recall that the goal here is to generate an outcome for each subject such that the overall synthetic data is more comparable with the original data set. We assume that the outcome variable $Y_i \in \{1, 0\}$ of each subject $i$ follows binomial distribution. Moreover, we model the probability of being a case is a linear function of the risk score $S_i = X_i^\top \hat{\beta}_{pre}$. Formally, we assume $E[Y_i|S_i] = M + aS_i$ for some $a \in \mathbb{R}$. Here $M \in (0, 1)$ is the proportion of case-subjects in the GWAS study and $E[Y_i|S_i]$ denotes the conditional expectation of random variable $Y_i$ given the risk score. For $1 : 1$

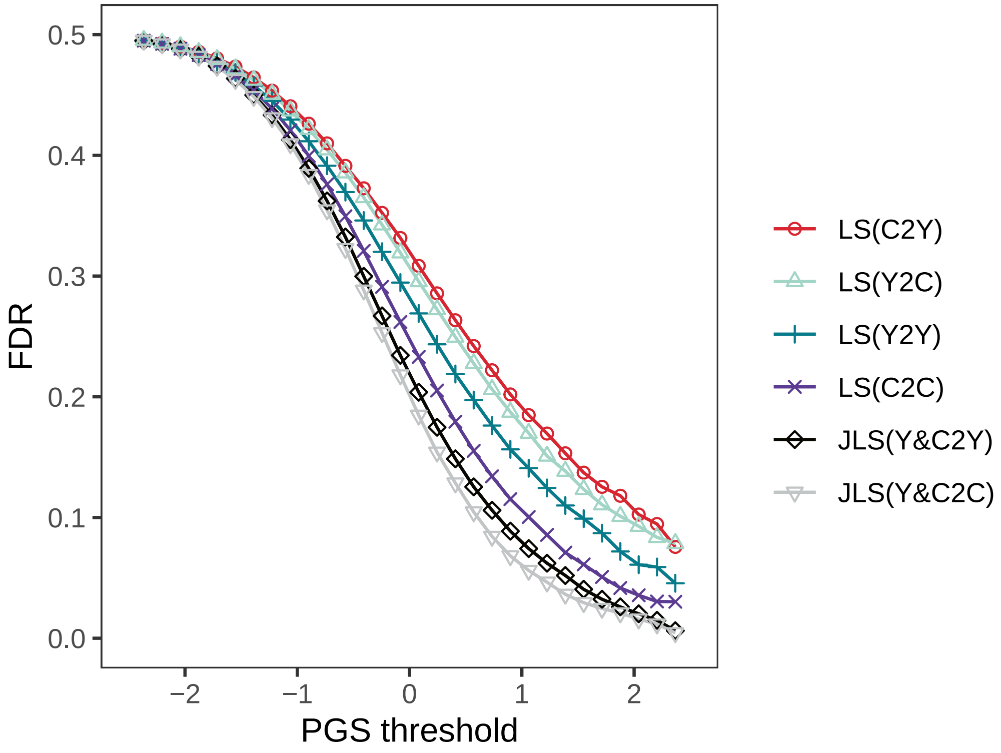


Figure S4: FDR,equal sample size setting.

case-control study, $M = 1/2$. When the expectation of risk score is zero $ES_i = 0$ we have $EY_i = M$. This means the generated data would have essentially the same study design as the original data.

Once we estimate the calibration slope $a$, we will have a probability for subject $i$ being a case sample and we can generate our synthetic data. We state that $a$ can be calculated using the following formula

$$a = \frac{\sqrt{n} c_M r^\top \hat{\beta}_{pre}}{\sum_{i=1}^n s_i^2}, \tag{2}$$

where $c_M^2 = M(1-M)$, $r \in \mathbb{R}^p$ is the Pearson-correlation vector between gene and the outcome $Y$ that is obtained from GWAS summary statistics.

We present the derivation of the calibration formula (2) below.

To estimate $a$ from data, we formulate $(X_i, S_i, Y_i)$ as independent and identically distributed sample from some common distribution $(X, S, Y)$. The variance of $Y$ is $M(1-M)$ (note that $Y$ is a binomial random variable with mean $M$). Denote $c_M^2 = M(1-M)$, we have:

$$\begin{aligned} Y &= E[Y|S] + Y - E[Y|S] \\ \Rightarrow Y - E[Y] &= aS + Y - E[Y|S] \\ \Rightarrow var(Y)^{-1/2}(Y - EY) &= c_M^{-1}(aS + Y - E[Y|S]) \\ \Rightarrow var(Y)^{-1/2}(Y - EY)S &= c_M^{-1}(aS^2 + SY - SE[Y|S]) \end{aligned} \tag{3}$$

Recall that $S = X^\top \hat{\beta}_{pre}$, we have

$$var(Y)^{-1/2} E[(Y - EY)X^\top]\hat{\beta}_{pre} = c_M^{-1} a E[S^2] \tag{4}$$

Since the genotype matrices are normalized to have zero mean and unit norm for each column, the variance of each dimension of $X$ is approximately $n^{-1}$. So approximately we have,

$$n^{-1/2} var(Y)^{-1/2} E[(Y - EY)(X - E[X])^\top]\hat{\beta}_{pre} = n^{-1/2} c_M^{-1} a E[S^2] \tag{5}$$

Since $n^{-1/2} var(Y)^{-1/2} E[(Y - EY)(X - E[X])]$ is the correlation coefficient between each SNP and the phenotype, we can estimate it using the GWAS summary statistic $r$. So we have

$$a = \sqrt{n} c_M r^\top \hat{\beta}_{pre} (E[S^2])^{-1} \sim \sqrt{n} c_M r^\top \hat{\beta}_{pre} (n^{-1} \sum_{i=1}^n S_i^2)^{-1}, \tag{6}$$

which is the proposed formula.

**Algorithm 1** Parameter Tuning with Synthetic Data.

**input:** GWAS summary statistics $\boldsymbol{r}_{pop1}, \boldsymbol{r}_{pop2} \in \mathbb{R}^p$, genotype matrices $\boldsymbol{X}_{pop1}, \boldsymbol{X}_{pop2}$; the proportion of case samples in the GWAS study, $M_{pop1}, M_{pop2}$ .

**Step 1:** Fit Joint-Lassosum models using $r_{pop1}, r_{pop2}$ under multiple candidate hyperparameters $(\lambda_1, \gamma_1), ..., (\lambda_H, \gamma_H)$. Each of the $H$ Joint-Lassosum models gives one regression coefficient vector $\hat{\beta}_1, ..., \hat{\beta}_H$. Choose one of them to be $\hat{\beta}_{pre}$;

**Step 2:** Generate Synthetic Data:

**for** *Each population* $i \in \{1, 2\}$ **do**

  **for** *Each* $j \in 1, ...,$ *row number of* $X_{popi}$ **do**

    Obtain genotype data $x \in \mathbb{R}^p$ of one subject, which we assume is the $j$-th row of $X_{popi}$;

    Calculate PGS $s_{ij} \leftarrow x\hat{\beta}_{pre}$ for this subject;

  **end**

  Calculate the calibration slope $a_i$ given by formula (2);

  **for** *Each* $j \in 1, ...,$ *row number of* $X_{popi}$ **do**

    Draw a random number $Y^*_{ij} \in \{0, 1\}$ from a Binomial distribution with mean $= M_i^{-1} + a_i s_{ij}$;

  **end**

  Split the subjects into a training and a testing set. Each subject has a genotype vector and a synthetic label $Y^*_{ij}$. (SS)

  Perform GWAS to get gene-phenotype correlation $r^*_{popi}$ using $X_{popi}$ and $\{Y^*_{ij}\}$.

**end**

**Step 3:** Fit Joint-Lassosum using the same $H$ hyperparameters as in **Step (1)**, but replace the summary statistics $r_{popi}$ by $r^*_{popi}$. Obtain regression coefficients $\hat{\beta}^*_1, ..., \hat{\beta}^*_H$.

**Step 4:** Evaluate the AUCs of $\hat{\beta}^*_1, ..., \hat{\beta}^*_H$ using the testing samples created in step (SS) above. Assume the $h$-th model $\hat{\beta}^*_h$ has the highest AUC.

**output:** return $(\lambda_h, \gamma_h)$, the selected hyperparameter, and its corresponding Joint-Lassosum model $\hat{\beta}_h$
